# The Site Frequency Spectrum under Finite and Time-Varying Mutation Rates

**DOI:** 10.1101/375907

**Authors:** Andrew Melfi, Divakar Viswanath

## Abstract

The diversity in genomes is due to the accumulation of mutations and the site frequency spectrum (SFS) is a popular statistic for summarizing genomic data. The current coalescent algorithm for calculating the SFS for a given demography assumes the *μ* → 0 limit, where *μ* is the mutation probability (or rate) per base pair per generation. The algorithm is applicable when *μN*, *N* being the haploid population size, is negligible. We derive a coalescent based algorithm for calculating the SFS that allows the mutation rate *μ*(*t*) as well as the population size *N*(*t*) to vary arbitrarily as a function of time. That algorithm shows that the probability of two mutations in the genealogy becomes noticeable already for *μ* = 10^-8^ for samples of *n* = 10^5^ haploid human genomes and increases rapidly with *μ*. Our algorithm calculates the SFS under the assumption of a single mutation in the genealogy, and the part of the SFS due to a single mutation depends only mildly on the finiteness of *μ*. However, the dependence of the SFS on variation in *μ* can be substantial for even *n* = 100 samples. In addition, increasing and decreasing mutation rates alter the SFS in different ways and to different extents.

## Introduction

Haldane [1947] noted that the gene for hemophilia appeared to mutate more rapidly in men than in women (also see [Rahbari et al., 2016]). Since then, a great deal of evidence has accumulated that the mutation rate varies across the human genome. CpG junctions are well-known to have particularly high mutation rates [Ehrlich and Wang, 1981]. Some of the variation in the mutation rate across the human genome appears to be related to correlations between single nucleotide polymorphisms found in humans as well as other nearby primates [Harpak et al., 2016, Hodgkinson et al., 2009, Hodgkinson and Eyre-Walker, 2011, Johnson and Hellmann, 2011]. A variety of statistical models of the variation of the mutation rate have been proposed and examined [Aggarwala and Voight, 2016, Carlson et al., 2018, Eyre-Walker and Eyre-Walker, 2014, Hodgkinson and Eyre-Walker, 2011, Michaelson et al., 2012]. The mutation rate can vary even between human populations [Harris, 2015, Mathieson and Reich, 2017, Narasimhan et al., 2017].

The site frequency spectrum (SFS) is commonly used to summarize the effect of mutations across the genome. For a haploid sample of size *n*, the SFS consists of the probability that *j* of the samples carry the mutant allele, for *j* = 1,…, *n* – 1, at a polymorphic site. If the mutation rate itself varies widely across the genome, it is essential to know how the mutation rate affects the SFS. In this article, we derive an algorithm to calculate the SFS with mutation rates as well as population sizes allowed to vary in an arbitrary manner. The mutation rate is assumed to be *μ*(*t*) per base pair per generation at time *t* and the haploid population size is assumed to be *N*(*t*). The algorithm relies on the coalescent approximation to genealogies [Durrett, 2008]. In particular, *n* samples are assumed to experience a binary merger according to a Poisson process of rate *n*(*n* – 1)/2*N*(*t*). The samples are hit with mutations by an independent Poisson process of rate *μ*(*t*)*n*.

An algorithm for calculating the SFS assuming *μN* to be negligible and *μ* to be constant in time is due to Polanski and Kimmel [2003]. The Polanski-Kimmel algorithm, which relies on the earlier work of Griffiths and Tavaré [1998] as well as Polanski et al. [2003], is based on the internal structure of the coalescent genealogy. In particular, the algorithm relies on the expected branch length of the genealogy with exactly *b* descendants for *b* = 1,…, *n* – 1.

Our algorithm allows *μ*(*t*) to be finite and varying in time and is also based on the coalescent approximation. However, it pays no attention to the internal structure of the genealogy. The algorithm is more Markovian in spirit and is partly based on the ideas in our earlier analytic work [Melfi and Viswanath, 2018a,b].

Harpak et al. [2016] have presented data analysis showing that samples of size *n* ≈ 10^5^ have experienced more than one mutation at several polymorphic sites with *μ* ∈ [10^-9^, 10^-7^]. Our algorithm can calculate the probability that a polymorphic site has experienced more than one mutation exactly. Using the demography inferred by Harpak et al. [2016], we precisely delineate the probability of more than one mutation in the genealogy.

When *n* and *μ* are large enough that a polymorphic site has been hit with either one, two, or more mutations, the SFS is a mixture of the SFS due to a single mutation and the SFS due to two or more mutations. Our calculations imply that the effects described by Harpak et al. [2016], such as the change in the profile of rare alleles at sites of higher *μ*, are mostly due to two or more mutations in the genealogy, which is in agreement with their conclusions.

Beginning with the work of Hwang and Green [2004], a number of authors have questioned the constancy of *μ*(*t*) with respect to *t* [Kim et al., 2006, Moorjani et al., 2016b,a, Narasimhan et al., 2017, Scally and Durbin, 2012]. The germ line mutation rate is known to depend on the number of cell divisions experienced by the germ line [Gao et al., 2016, Amster and Sella, 2016]. The number of cell divisions in the germ line is greater in the human male than the human female, and in the male it increases with age. The dependence on the number of cell divisions could be due to errors during either genome replication or DNA repair [Gao et al., 2016], and some of the de novo mutations are shared between siblings in a mosaic pattern [Rahbari et al., 2016]. We use our algorithm to illustrate how increasing and decreasing mutation rates alter the SFS.

The coalescent and the diffusion equation often provide alternative routes to the same results. Accordingly, the SFS can be computed using the diffusion equation [Griffiths, 2003, Gutenkunst et al., 2009, Sawyer and Hartl, 1992]. For the possibility of handling varying mutation rates using the diffusion equation, see [Steinrücken et al., 2016, Živković et al., 2015]. In the diffusion approach, the transition probabilities are first obtained and the SFS is computed using the transition probabilities. The sample size *n* enters only during the latter step. Therefore, it may appear as if the diffusion equation can calculate SFS for even large *n* with not much more trouble than for small *n*. However, for large *n*, the transition probabilities have to be calculated more accurately and with greater resolution.

### Calculating the SFS under Finite and Varying Mutation Rates

Suppose the population size *N*(*t*) ≡ *N* and mutation rate *μ*(*t*) ≡ *μ* are both constant. If a site is polymorphic, the probability that *j* out of *n* samples carry the mutant allele converges to

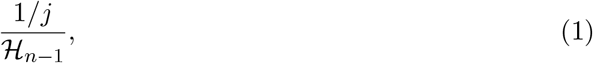

where 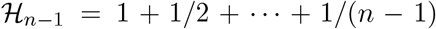, in the limit *μN* → 0 [Durrett, 2008]. Before presenting the full algorithm, we treat the case where *N* and *μ* are both constant but without the assumption of *μN* being negligible. The extension of the classical result (1) that we derive brings out some of the main ideas in a relatively transparent way.

If the coalescent genealogy is sectioned at some fixed time in the past, we will refer to the lineages present at that time in the past as ancestral samples, following our earlier usage [Melfi and Viswanath, 2018a,b]. Suppose the number of ancestral samples is *k*. Because the Poisson process of rate *k*(*k* – 1)/2*N* that produces a binary merger in the ancestral sample and the Poisson process of rate *μk* that hits the sample with a mutation are independent, the probability that the next event in the genealogy is a binary merger is

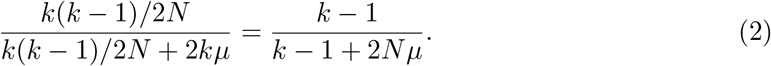

Correspondingly, the probability that the next event is a mutation is

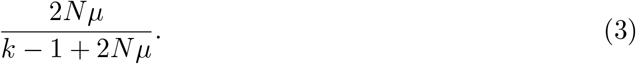

It follows that the probability *q*_0_(*k*) that *k* samples coalesce without being hit by a mutation is

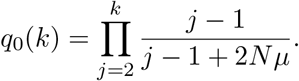

In more detail, for ancestral sample sizes *j* = 2,…, *k*, a binary merger must precede a mutation, which occurs with a probability given by (2) (with *k* ← *j*) for each *j* = 2,…,*k*.

Similarly, the probability *q*_1_(*k*) that a sample of size *k* coalesces after experiencing exactly one mutation is

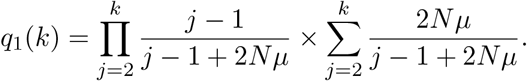

In more detail, the probability that a sample of size *k* coalesces after experiencing exactly one mutation when the ancestral sample size is *j* but with no other mutations in the genealogy is

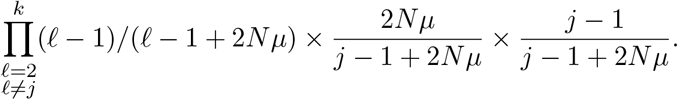

The first factor occurs because the first event to hit an ancestral sample of size *ℓ* ≠ *j* is a binary merger. The second factor occurs because the first event experienced by an ancestral sample of size *j* must be a mutation (whose probability is given by (3) with *k* ← *j*) and the third factor because the sample of size *j* then experiences a binary merger (whose probability is given by (2) with *k* ← *j*). The formula for *q*_1_(*k*) is obtained by summing over *j* = 2,…,*k*.

The condition or event that *n* samples coalesce with exactly one mutation in the genealogy will be denoted by 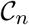. The definition of 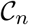 will be changed slightly after we introduce the concept of an ancestral lens. Conditioned on 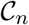, the probability that a mutation event occurs in the genealogy when the sample size is *k* is given by

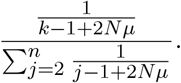

Using the coalescent propagators derived earlier [Griffiths and Tavaré, 1998, Melfi and Viswanath, 2018b], we may deduce the probability of *j* mutants in the sample under the condition 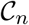 to be

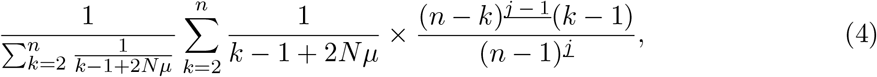

where 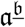 denotes the falling power 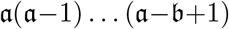. The SFS (1) is obtained by substituting *μ* = 0 in this formula.

There is no obvious way to evaluate this formula for *j* = 1,…,*n* – 1 in 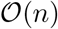 arithmetic operations for *μ* > 0, although the *μ* = 0 case given by (1) can be evaluated in 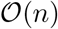 arithmetic operations. The formula (4) for the SFS under the condition 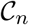 can be cast in the form of a recurrence and the SFS evaluated using 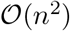 operations [Melfi and Viswanath, 2018a].

This derivation of the SFS conditioned on 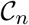 for constant *N* and *μ* relies on *q*_0_(*k*), the probability that *k* samples coalesce with zero mutations, and *q*_1_(*k*), the probability that *k* samples coalesce with exactly one mutation in their genealogy. The algorithm with varying *N*(*t*) and *μ*(*t*) uses a similar approach. The more general algorithm relies on the concept of an ancestral lens, to which we now turn, to reduce the number of arithmetic operations.

### Ancestral and calendar lenses

We denote ancestral time by *τ* (generations), with the current epoch being *τ* = 0. If the number of samples is *n*, let *p*(*k*, *τ*) be the probability that the number of ancestral samples at time *τ* is *k*. The ancestral lens is defined as the set of all (*k, τ*) such that *p(k,τ*) ≥ *ϵ_lens_*. The tolerance *ϵ_lens_* is taken to be 10^-40^. All sample sizes outside the ancestral lens have a negligible probability and may be ignored without affecting any calculations for the current sample at *τ* = 0. Significant savings may be realized by confining calculations to the ancestral lens.

The probability that the ancestral sample is of size *k* at time *τ* and undergoes a binary merger in the interval [*τ, τ* + *dτ*] is

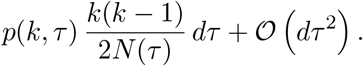

Likewise, the probability that the ancestral sample is of size *k* + 1 and undergoes a binary merger in [*τ, τ* + *dτ*] is

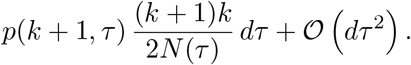

Therefore,

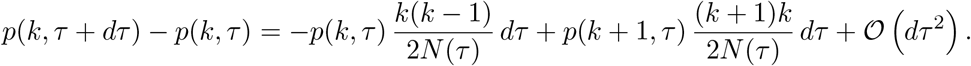

In the limit *dτ* → 0, we obtain the differential equation

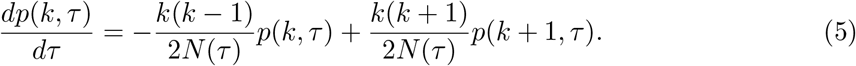

To calculate the ancestral lens, this differential equation is initialized with *p*(*n*, 0) = 1 and *p*(*k*, 0) = 0 for *k* ≠ *n*. The numerical method used for computing *k_min_*(*τ*) and *k_max_*(*τ*) such that *p*(*k, τ*) < *β_lens_* for *k* ∉ [*k_min_*(*τ*),*k_max_*(*τ*)] is described in the appendix. The functions *k_min_*(*τ*) and *k_max_*(*τ*) are the boundaries of the ancestral lens.

Figure 1 shows the ancestral lens for a demographic model. The considerable savings realized by confining calculations to the ancestral lens are obvious from that figure. We work with three demographic models:

**Fig. 1:**
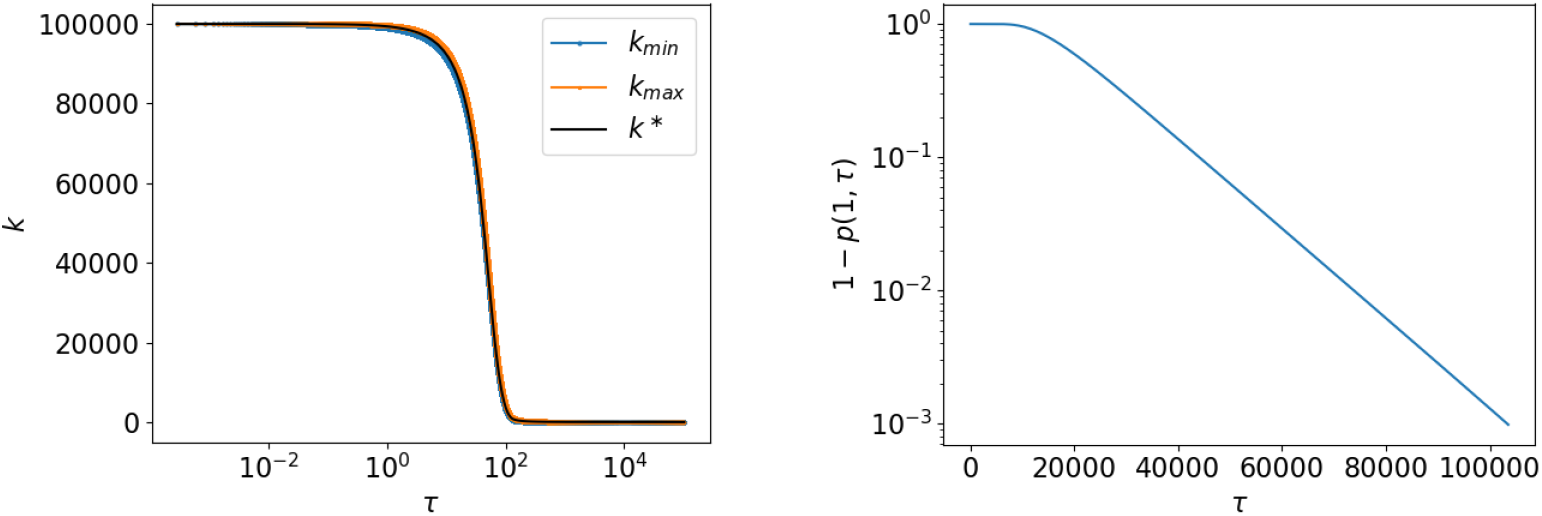
The ancestral lens with *n* = 10^5^ and demographic model 2. The second plot shows the probability of (non)coalescence as a function of ancestral time *τ*.

- Demographic model 0: *N* is assumed to be constant and equal to 2 × 10^4^.
- Demographic model 1: *N*(*τ*) = *N*_0*e*_^−*rτ*^, with *N*_0_ = 8×10^6^ and *r* = 0.017, for *τ* ∈ [0,367.8] and *N*(*τ*) = *N*_0*e*_^-*r*367.8^ ≈ 15403 for *τ* ≥ 367.8. This model is from [Nelson et al., 2012].
- Demographic model 2: *N*(*τ*) = *N*_0*e*_-^*rτ*^, with *N*_0_ = 8 × 10^6^ and *r* = 0.0538, for *τ* ∈ [0,119.47] and *N*(*τ*) = *N*_0*e*_^-*r*119.47^ ≈ 12932 for *τ* ≥ 119.47. This model is from [Harpak et al., 2016].

Figure 1 uses model 2. In that figure, the stopping time *τ* = *T* for the ancestral lens is determined using the criterion 1 – *p*(1, *T*) ≥ 10^-3^, which ensures that the entire sample has coalesced with probability greater than 99.9%. Such a criterion makes *T* very large, however, as shown in the figure. It is better to stop the lens when *μ* and *N* have become constant. The entire section of the lens at the stopping time *T* can be initialized using exact formulas for *q*_0_, *q*_1_, and the SFS.

The computation of the ancestral lens is forward in ancestral time *τ*. However, the computation of *q*_0_, *q*_1_, and the SFS march forward in calendar time. Therefore, the ancestral lens must be flipped to a calendar time lens as shown in Figure 2. In both Figures 1 and 2, *k*\* is the most likely size of the ancestral sample. We take the stopping time *τ* = *T* to be the origin *t* = 0 of calendar time so that the current epoch is *t* = *T*.

**Fig. 2:**
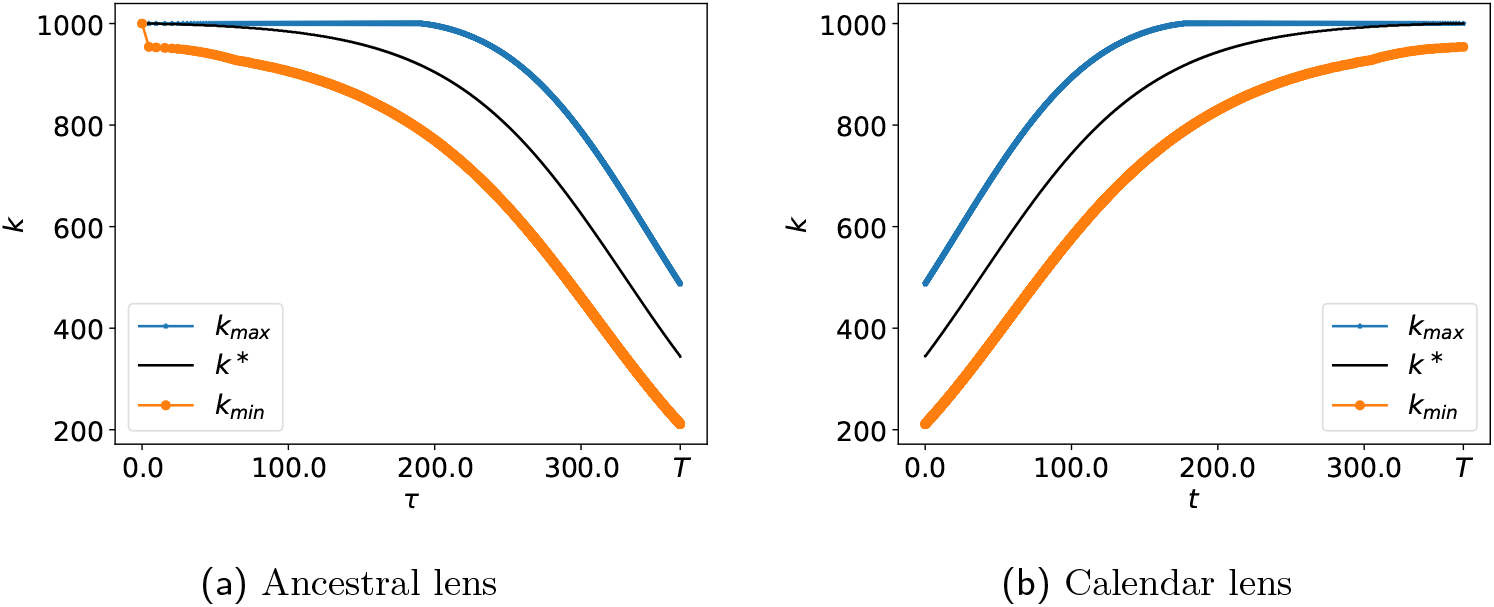
The ancestral and calendar lenses for *n* = 1000 samples under demographic model 1.

### Computation of *q*_0_(*k,t*) and *q*_1_(*k,t*)

The probability that *k* samples at calendar time *t*, *t* ∈ [0, *T*], coalesce within the lens or exit the lens without being hit by a mutation is denoted *q*_0_(*k,t*). The probability of exiting the lens is very low and we regard any exit from the lens as equivalent to coalescence. Any loss of accuracy will be below the level of machine precision. Thus, *q*_0_(*k,t*) = 1 for *k* ∉ [*k_min_* (*t*), *k_max_*(*t*)]. In addition, *q*_0_(1,*t*) = 1.

Suppose there are *k* ancestral samples at time *t*. If we go back to time *t* – *dt* and ignore *dt*^2^ terms, we have the following possibilities:

- The sample is hit with a mutation with probability *μ*(*t*)*k dt*.
- There is a binary merger with probability *k*(*k* – 1) *dt*/2*N*(*t*).
- There is neither a mutation nor a binary merger with probability 1 – *μ*(*t*)*k dt* – *k*(*k* – 1) *dt*/2*N*(*t*).

Because we are ignoring *dt*^2^ terms, these three possibilities are disjoint and exhaustive.

We have

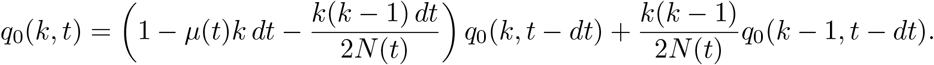

If the *k* samples are neither hit by a mutation nor experience a binary merger in [*t* – *dt,t*], we require that *k* samples at time *t* – *dt* coalesce (or exit the lens) without a mutation. If the samples experience a binary merger, we require that *k* – 1 samples at time *t* – *dt* coalesce (or exit the lens) without a mutation. In the *dt* → 0 limit, we have the following differential equation:

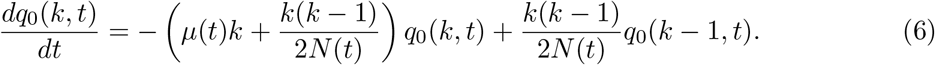

The differential equation is solved for *t* ∈ [0, *T*] as an initial value problem. For *k*(0) ≤ *k* ≤ *k_max_*(0), *q*_0_(*k*, 0) is initialized using the exact formula for *q*_0_(*k*) when *μ* and *N* are constant. After solving the initial value problem, we obtain *q*_0_(*n,T*), which is the probability that *n* samples at the current epoch coalesce (or exit the lens) without being hit by a mutation.

Similarly, let *q*_1_(*k,t*) be the probability that *k* ancestral samples at time *t* coalesce (or exit the lens) while being hit by exactly one mutation. In this case, the same three disjoint and exhaustive possibilities imply

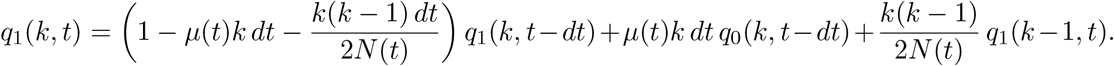

If there is neither a binary merger nor a mutation in [*t* – *dt*, *t*], we require that *k* samples at time *t* – *dt* coalesce (or exit the lens) with exactly one mutation. If there is a mutation event, we require that *k* samples at time *t* – *dt* coalesce (or exit the lens) without suffering a mutation. Finally, if there is binary merger, we require that *k* – 1 samples at time *t* coalesce (or exit the lens) while suffering exactly one mutation.

In the limit *dt* → 0, we have the differential equation

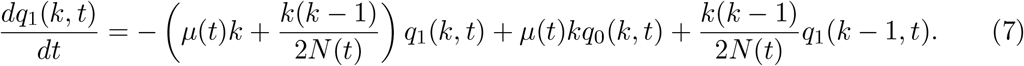

This differential equation too is solved as an initial value problem from *t* = 0 to *t* = *T*. For *k* ∈ [*k_min_*(0), *k_max_*(0)], *q*_1_(*k*, 0) is initialized using the exact formula for *q*_1_(*k*) with constant *μ* and *N*.

If *k* ∉ [*k_min_*(*t*), *k_max_* (*t*)], then *q*_1_(*k,t*) = 0. In addition, *q*_1_(1,*t*) =0 for *t* ∈ [0, *T*].

If *q*_2_(*k,t*) is the probability that *k* ancestral samples at time *t* coalesce with exactly 2 mutations, *q*_2_(*k,t*) satisfies an analogous differential equation:

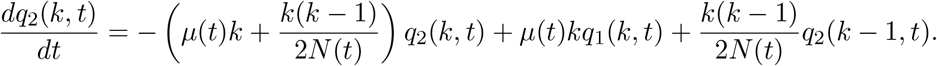

If *N* and *μ* are constant

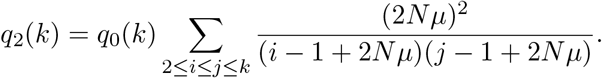

The initial value problem for *q*_2_(*k,t*) may be initialized using *q*_2_(*k*, 0) = *q*_2_(*k*) with *N* = *N*(0) and *μ* = *μ*(0).

### Computation of the SFS under the condition 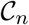

Suppose the probability that *j* samples out of *n* are mutants at a polymorphic site is *p_j_*. We may then represent the SFS using a generating function as 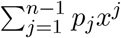. The generating function representation will be used presently to define the extension operator and to derive a differential equation for the SFS.

Suppose the generating function is denoted *S* and that *n* + 1 samples are related to the *n* samples via a binary merger. The probability that *j* out of *n* + 1 samples are mutants is given by

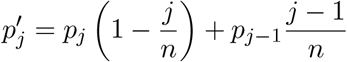

for *j* = 2,…,*n* – 1, by 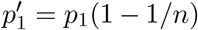 for *j* = 1, and 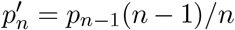 for *j* = *n*. Each of the *n* parents of the sample of size *n* + 1 is equally likely to split into two. Therefore, to end up with *j* mutants among *n* + 1 samples, we must have either *j* mutants among *n* samples with one of the non-mutants splitting (which occurs with probability (1 – *j/n*)) or *j* – 1 mutants among *n* samples with one of the mutants splitting (which occurs with probability (*j* – 1)/*n*). The extension operator 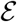 is defined by

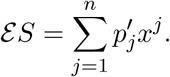

The extension operator 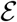 applied to the SFS *S* results in the SFS of a sample of size greater by one that is assumed to be related to the parental sample via a single binary merger.

Suppose *k* ∈ [*k_min_*(*t*), *k_max_*(*t*)]. Under the condition 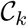 that those *k* ancestral samples coalesce (or exit the lens) with exactly one mutation, the *k* samples are hit with a mutation in the time interval [*t* – *dt*, *t*] with a probability equal to

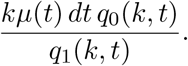

Thus, under the condition 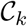, the *k* samples are hit with a mutation at the rate

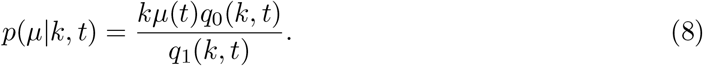

Similarly, the *k* samples experience a binary merger with the conditional rate given by

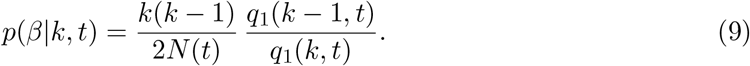

It must be noted that *p*(*μ*|*k,t*) and *p*(*β*|*k,t*) are rates and not probabilities.

Under the condition 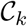, there are three disjoint and exhaustive possibilities (with *dt*^2^ terms ignored) for *k* ancestral samples in the time interval [*t* – *dt, t*].

- The *k* ancestral samples are hit with a mutation with probability *p*(*μ*|*k,t*) *dt*.
- The samples experience a binary merger with probability *p*(*β*|*k,t*) *dt*.
- There is no event with probability (1 – *p*(*μ*|*k,t*) – *p*(*β*|*k,t*)) *dt*.

Let *S*(*k, t*), with *t* ∈ [0, *T*] and *k* ∈ [*k_min_*(*t*),*k_max_*(*t*)], denote the SFS of *k* ancestral samples at time *t* under the condition 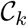. The three disjoint possibilities listed above imply that

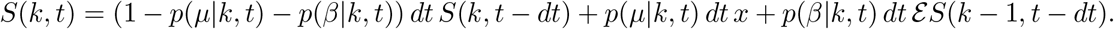

The middle term corresponds to a single mutant arising because of a mutation occurring in the interval [*t* – *dt, t*]. The last term corresponds to a binary merger.

In the limit *dt* → 0, we obtain the differential equation

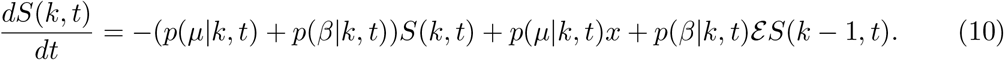

This differential equation is solved as an initial value problem from *t* = 0 to *t* = *T*. The SFS *S*(*k*, 0) for *k* ∈ [*k_min_*(0), *k_max_*(0)] is initialized using the exact formula when *μ* and *N* are both constant. For *k* ∉ [*k_min_*(*t*),*k_max_*(*t*)], *S*(*k,t*) = 0. In addition, *S*(1,*t*) = 0. The numerical solution of this differential equation is described in the Appendix. A computer program implementing the algorithm may be obtained from github.com/divakarvi/18-varymu.

### SFS under the Wright-Fisher model

The resulting limit is the zero mutation limit with varying mutation rate. An algorithm applicable to this limit is described in the Appendix.

SFS under the Wright-Fisher model

The Wright-Fisher model differs from the coalescent in that all mergers under the coalescent are binary mergers, whereas the Wright-Fisher model allows multiple mergers as well as simultaneous mergers [Durrett, 2008]. Harpak et al. [2016] considered whether the SFS computed assuming every merger to be a binary merger is reliable.

The formal truncation error in passing from the Wright-Fisher model to the coalescent is *n*^2^/*N* [Durrett, 2008]. However, the leading truncation term in the SFS is only *n*/*N* and in fact the total variation distance in the SFS is only around 1% for *n* = *N* = 20,000 [Melfi and Viswanath, 2018b]. Furthermore, in demographic models such as 1 and 2 characterized by recent exponential growth, the total variation distance may be expected to be even lower. Assuming every merger to be a binary merger has little effect on the SFS.

Nevertheless, in the Appendix, we show how to compute the SFS with varying *μ*(*t*) and *N*(*t*) under the Wright-Fisher model. An advantage of the Wright-Fisher model is that it is already discrete. We also remark that computations with the Wright-Fisher model may lend themselves to better optimization with the use of suitable asymptotic formulas.

### Effect of Finite Mutation Rate on the SFS

Harpak et al. [2016] have presented evidence that the infinite sites model is violated for samples of around *n* = 10^5^ haploid human genomes. The infinite sites models assumes that every new mutation in the genealogy occurs at a different site.

To understand when and how violations of the infinite sites model set in, we may first look at the quantity

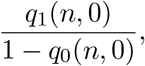

which is the probability that there is exactly one mutation in the genealogy given that there are one or more mutations in the genealogy of the *n* samples. In Figure 3, we use *q*_1_/(1 – *q*_0_) as a surrogate for the probability that a site is polymorphic as a result of a single mutation. The two probabilities will be close but are not exactly the same. They differ slightly because of the small probability that two mutations may occur in the same lineage in the genealogy and cancel each other. Thus, a sample with two or mutations in its genealogy may not be polymorphic.

**Fig. 3:**
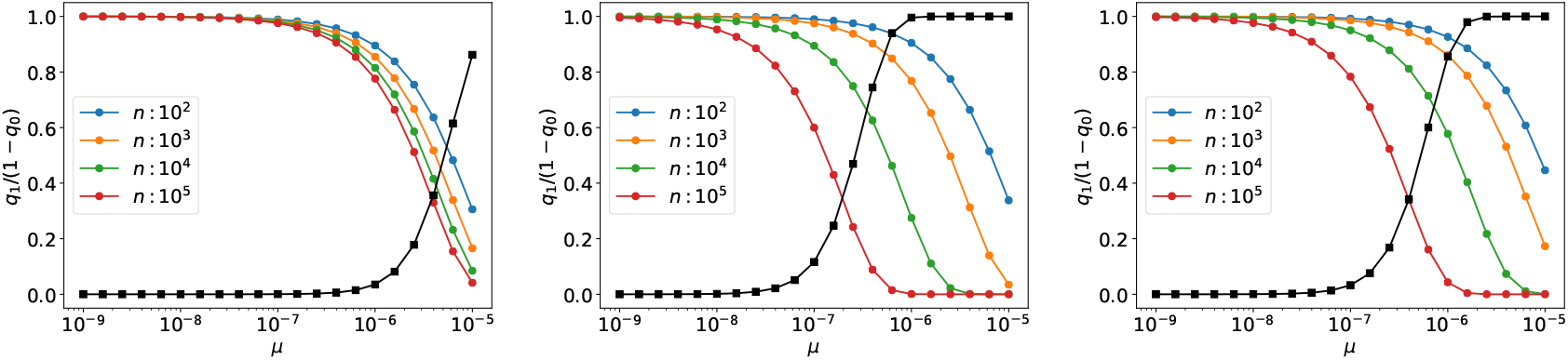
Probability that a site is polymorphic as a result of a single mutation in the genealogy for demographic models 0, 1, 2 and various sample sizes. In all three plots, the black squares plot the probability for *n* = 10^5^samples that a site hit with a mutation has been hit with three or more mutations.

In Figure 3, we have graphed the probability of a single mutation at a polymorphic site for demographic model 0 which assumes *N* ≡ 2 × 10^4^. Sample sizes of *n* = 10^5^ would not make sense in the Wright-Fisher interpretation of that model. However, in the coalescent *N* is only a parameter to control rates of binary mergers.

The probability of a single mutation at a polymorphic site is the highest in demographic model 0. That appears to be because binary mergers are initially fastest in demographic model 0.

Both demographic models 1 and 2 assume exponentials that persist for more than a 100 generations. The population explosion slows down binary mergers and as a result the probability of double mutations is higher in demographic models 1 and 2. The exponential persists over a greater interval of time in model 1 and model 1 shows a greater probability of double mutations than model 2. In both models 1 and 2, the probability of a multiple mutation is noticeable for even *μ* = 10^-8^ and *n* = 10^5^.

The probability that a site hit with a mutation has been hit with three or more mutations is given by

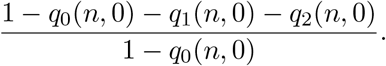

For demographic model 1, *n* = 10^5^, and *μ* = 10^-7^, the probability that a site that has been hit with a mutation has been hit with exactly one, exactly two, or three or more mutations is 60%, 28.4%, and 11.6%, respectively (see Figure 3). For demographic model 2, those probabilities are 78.3%, 18.4%, and 3.3%, respectively.

Harpak et al. [2016] have inferred a mutation rate of around 10^-7^ for CpG junctions. The SFS will be a mixture of the SFS due to a single mutation, the SFS due to two mutations, and the SFS due to three or more mutations in the genealogy. A sample being hit with three mutations at a site does not necessarily imply that the site shows four-fold polymorphism. Because transitions are more likely than transversions, the site may only be biallelic.

Suppose a sample carries multiple mutations at the same site in its genealogy. That does not imply that there is a haploid individual in the sample whose lineage has been hit with two mutations. The mutations may not be nested in the genealogical tree. If an ancestral lineage is hit with a mutation, the probability that the lineage is hit with yet another mutation may change [Nei, 1987].

In our analysis, we have assumed the probability to stay constant. Despite the limitations of that assumption, we may disentangle the effects of single and multiple mutations on the SFS. To do so, we turn to Figure 4. The total variation distance (or variation distance) between an

**Fig. 4:**
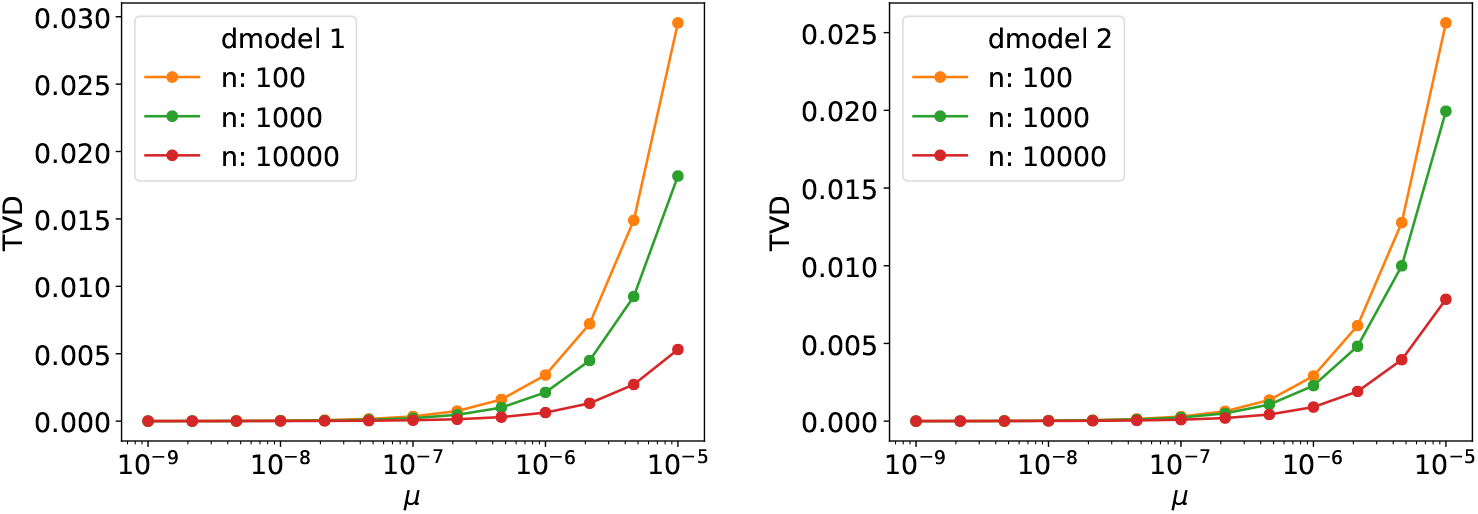
The total variation distance between the SFS at a given *μ* and at *μ* = 0 under the condition 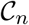.

SFS given by *p_j_* and an SFS given by 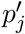 for *j* = 1,…,*n* – 1 is defined as

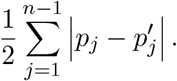

It is the right metric to use for comparison because it can be interpreted as the maximum difference between the probabilities of any possible event that is a subset of {1,…,*n* – 1} [Brémaud, 1999]. Because it can be interpreted as a probability, the variation distance can also be thought of as a percent.

Figure 4 shows that the variation distance of the SFS with *μ* finite and *μ* = 0 under the condition 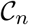 falls as the sample size *n* increases. The variation distance is negligible for *μ* = 10^-7^. Thus, the phenomena described by Harpak et al. [2016] are entirely due to multiple mutations in the genealogy.

In addition to the variation distance of the SFS between *μ* finite and *μ* = 0 being small, it is in a direction opposite to the total effect. Figure 5 shows that the effect of finite *μ* is to slightly increase the occurrence of rare alleles. However, in the overall SFS, rare alleles are depleted [Harpak et al., 2016]. Intuitively, we may understand why rare alleles are depleted by multiple mutations as follows. The probability of singletons (*j* = 1 mutants) begins to dominate in large samples as evident from Figure 5. When the genealogy carries two mutations, for example, there is a substantial chance that both mutations are transitions. In large samples and under demographic models with exponentially growing population, both mutations are likely to hit near the leaves of the genealogical tree. The mutations are unlikely to be nested. Therefore, the probability of a singleton is depleted.

**Fig. 5:**
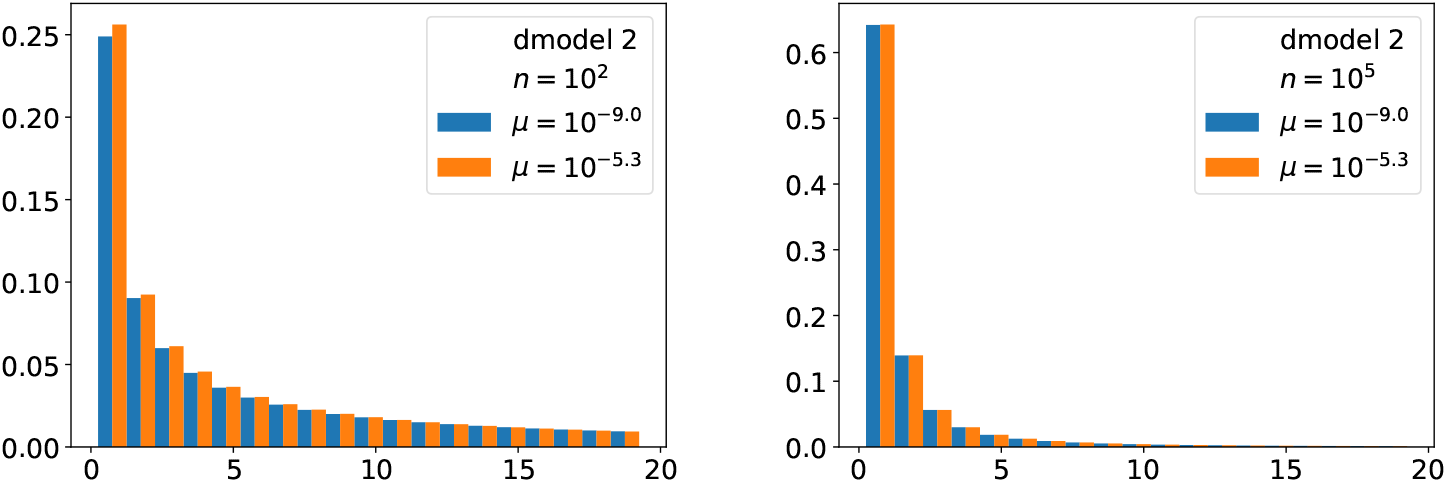
The effect of finite *μ* on the SFS (*j* < 20).

### The Effect of Varying Mutation Rate on the SFS

We assume the mutation rate to be given by

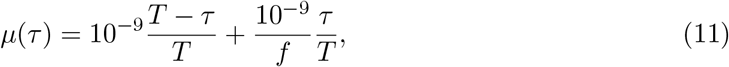

for *τ* ∈ [0, *T*], and *μ*(*τ*) = 10^-9^/*f* for *τ* ≥ *T*. The mutation rate is assumed to vary linearly in [0,*T*]. In addition, we assume *T* = 367.8 and *T* = 119.47 for demographic models 1 and 2, respectively. Because the historical variation of the mutation rate is only now being investigated, these assumptions are necessarily somewhat arbitrary. Genetic data is highly sensitive to *μ*, and there is no reason to assume anything faster than linear variation.

In the model for *μ*(*t*), the variation in *μ* sets in at *τ* = *T*. For demographic models 1 and 2, we have assumed *T* to be to the epoch at which exponential increase in population sets in.

In the model, *f* is the factor by which the mutation rate increases from *τ* = *T*, which is *T* generations in the past, to *τ* = 0, which is the current time. We have assumed *μ*(*t*) to be 10^-9^ at the current time because that is the most widely quoted number, although it has been questioned recently [Gao et al., 2016, Scally and Durbin, 2012].

Methods used to infer the mutation rate rely directly or indirectly on the pioneering work of Watterson [1975] on the number of segregating sites. For a recent example, see [Bhaskar et al., 2015]. Therefore, it is appropriate to begin by looking at the probability 1 – *q*_0_(*n*, 0) that a current sample of size *n* has been hit with a mutation at a site. The probability 1 – *q*_0_(*n*, 0) is close to but not exactly the same as the probability that a site is segregating. If a site is hit with multiple mutations, there is a small probability that it is not segregating.

Figure 6 shows the way the number of segregating sites (more precisely, the probability a site is hit with a mutation) varies as a function of *f*. A sample size of *n* = 100 appears more sensitive to variations in the mutation rate than larger samples, especially when *f* > 1 and *μ*(*t*) is increasing.

**Fig. 6:**
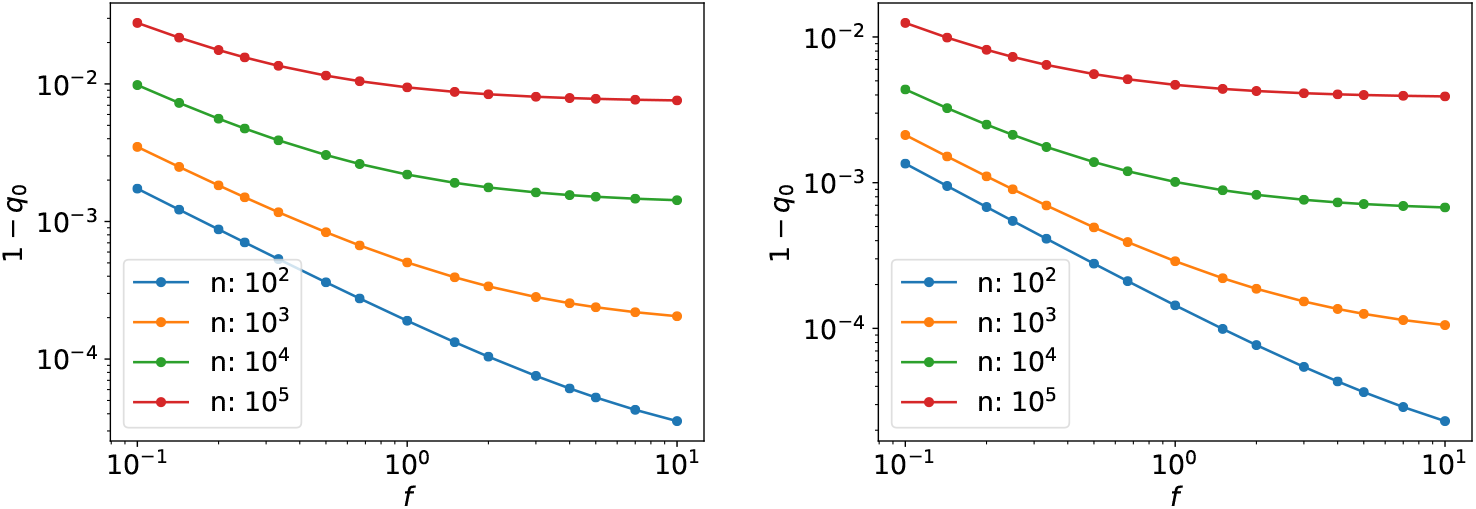
The probability that a site is segregating (more precisely, hit with a mutation) as a function of the factor *f* (see (11)) for demographic models 1 and 2, respectively.

Figure 7 shows the total variation distance of the SFS between *f* = *f* and *f* = 0 as a function of the parameter *f*. There is no easily discernible pattern in the plots. For the demographic model 1 and *n* = 100, the SFS changes by 6% and 3% when the mutation rate is assumed to double or halve, respectively. For demographic model 2, those numbers are 5% and 3%, respectively. The variation in *μ*(*t*) certainly has an effect on the SFS. Figure 8 shows that an increasing mutation rate augments the fraction of rare alleles, whereas a decreasing mutation rate depletes it.

**Fig. 7:**
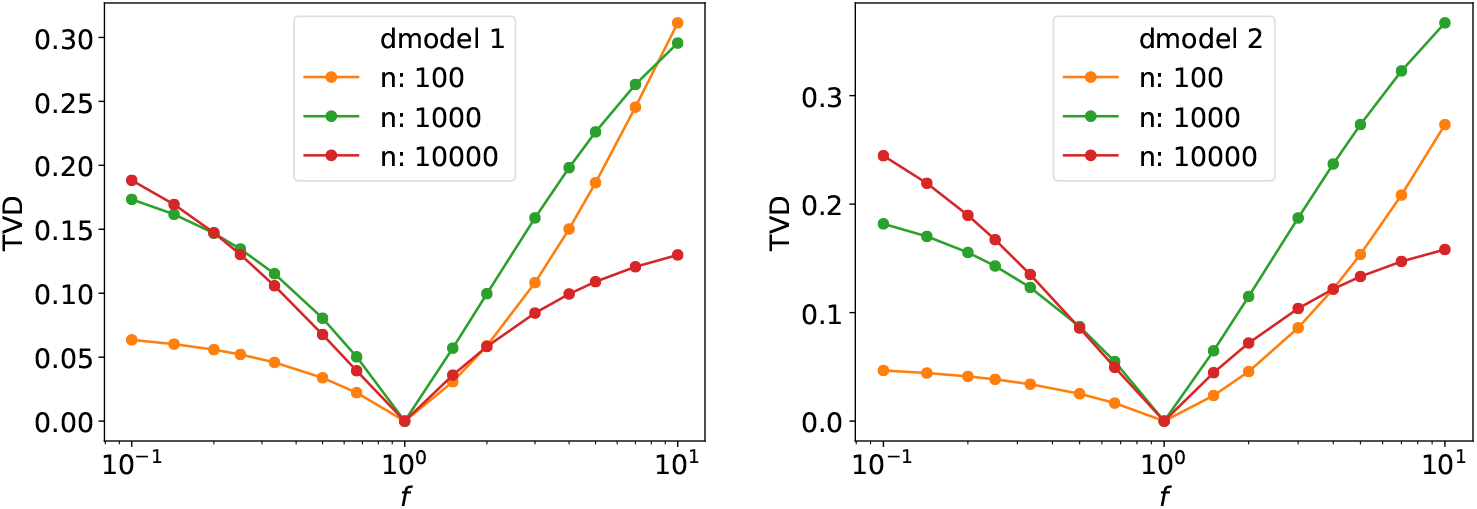
Total variation distance of the SFS for a given *f* (see (11)) from the SFS with *f* = 0, which implies a constant mutation rate. The parameter *f* controls the variation in mutation rate.

**Fig. 8:**
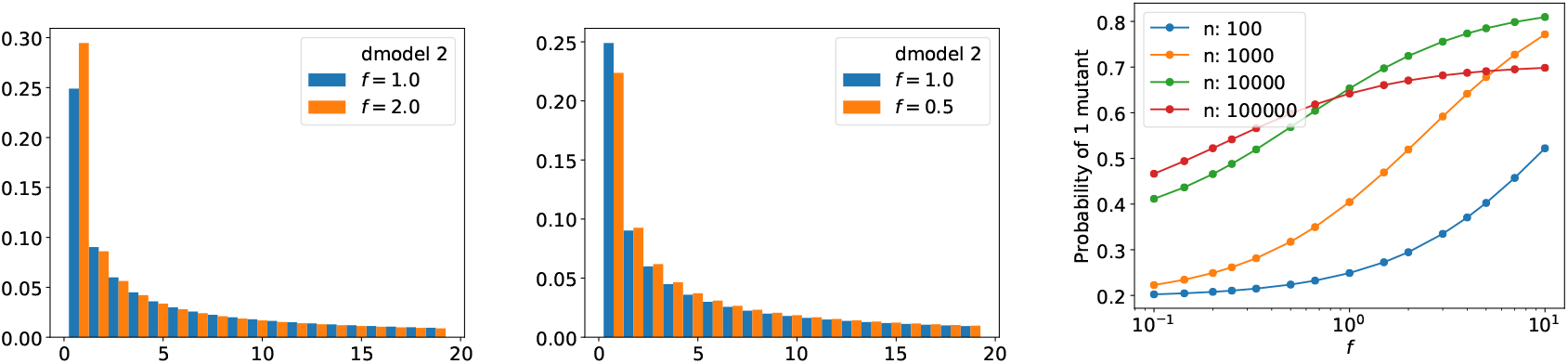
The effect of increasing and decreasing mutation rates on the SFS. The last plot uses demographic model 2 as well.

## Discussion

The Polanski-Kimmel algorithm for computing the SFS [Polanski and Kimmel, 2003] allows the population size *N*(*t*) to vary but assumes the mutation rate to be constant and negligible. We have presented an algorithm to compute the SFS that allows the mutation rate *μ*(*t*) to be finite and varying. The main innovation in our algorithm is to ignore the internal structure of the genealogical tree and instead take a more Markovian approach.

The algorithm uses a first pass (with ancestral time *τ* increasing) to calculate the ancestral lens. The ancestral lens is then flipped to the calendar time lens. In the second pass, the SFS as well as *q*_0_(*k,t*) and *q*_1_(*k,t*) are calculated with calendar time *t* increasing. The calculations solve a set of linear ordinary differential equations with time-varying coefficients.

The quantities *q*_0_(*k,t*) and *q*_1_(*k,t*) are the probabilities that a sample of size *k* coalesces without being hit by a mutation or after being hit by exactly one mutation, respectively. The algorithms for calculating *q*_0_ and *q*_1_ may be thought of as generalizations of the work of Watterson [1975] on the number of segregating sites. The generalization consists in allowing both *N*(*t*) and *μ*(*t*) to vary arbitrarily.

When the sample size is *n* ≈ 10^5^ and the mutation rate is *μ* ≈ 10^-7^, multiple mutations occur in human genealogies with appreciable probabilities. Algorithms for obtaining the SFS with two mutations have been applied to sample sizes of *n* = 100 [Jenkins et al., 2014]. In addition to reaching greater sample sizes, possibly while limiting calculations to rare alleles, there are yet other issues to be investigated. For example, once a site is hit with a mutation, the probability that it will be hit with yet another mutation will change. If the mutation probability is assumed to be constant, our calculations show that a polymorphic site may be hit with three mutations with a probability of around a few percent.

## Appendix

In the appendix, we explain how the ancestral lens and the SFS are computed numerically. We also give a version of the algorithm that allows *μ*(*t*) to vary and then takes the zero mutation limit. In addition, we show how to calculate the SFS for the Wright-Fisher model with varying mutation rate and population size.

### Implementation of the ancestral lens

The differential equation (5) is linear with non-constant coefficients. The decay rate *k*(*k* – 1)/2*N*(*τ*) is around 10^4^ for *k* ≈ 10^5^ and *N* ≈ 10^6^. Yet, the differential equations for *k* = 2,…,*n* cannot be considered stiff because the decay rate corresponds to the rate of binary mergers and must be resolved. Out of precaution, we used a 4th order BDF discretization [Hairer et al., 1991]. There is no additional cost to using an implicit method because the equations are linear. Initial time-steps were taken using the implicit midpoint rule. The formal order of accuracy is 3.

Integrating the differential equations for *k* = 2,…,*n* would be too expensive for large *n*. Instead, we restrict the integration to the ancestral lens even as it is being computed. Suppose the ancestral lens at *τ* is given by [*k_min_, k_max_*]. In the time step from *τ* to *τ* + *h*, we use *k* = max(*k_min_* – 1, 1),…, *k_max_*. After the time step, *k_min_*(*τ* + *h*) can be either *k_min_* or *k_min_* – 1. It is equal to *k_min_* – 1 if *p*(*k_min_* – 1, *τ* + *h*) < 10^-40^ and *k_min_* otherwise. Similarly, *k_max_*(*τ* + *h*) can be either *k_max_* – 1 or *k_max_*. It is equal to *k_max_* – 1 if *p*(*k_max_,τ* + *h*) < 10^-40^ and *k_max_* otherwise.

Particular care is necessary during the very first time step. The ancestral lens at *τ* = 0 is given by *k* ∈ [*n, n*] and consists of a single point. If the above strategy is followed, the ancestral lens can grow to at most *k* ∈ [*n* – 1, *n*] after the first time step. The tolerance of *ϵ_lens_* = 10^-40^ is so small that the ancestral lens will in fact be much wider. Failing to capture its width correctly in the first step will corrupt the entire computation. To capture the width of the ancestral lens correctly, the implicit midpoint rule is iterated until the width of the ancestral lens stabilizes. The way the ancestral lens grows during the very first step may be observed from Figure 2a.

Figure 2b shows an ancestral lens flipped to a calendar time lens. The flip is mostly straightforward except at the current epoch *t* = *T* or *τ* = 0. At *τ* = 0, the ancestral lens is given by *k* ∈ [*n, n*]. However, *k* ∈ [*n, n*] at *t* = *T* will not do because the differential equations (6), (7), and (10) utilize information regarding *k* – 1 samples. When an implicit BDF discretization is involved, narrowing the lens suddenly at *t* = *T* will create error in moving information from *n* – 1 samples to *n*. This problem is easily solved by taking the calendar time lens at *t* to be equal to the ancestral lens at *τ* = *T* – *t* + *h*, where *h* is the time step into *t*.

### Solution of the differential equations for *q*_0_, *q*_1_, and the SFS

Provided the calendar time lens is calculated carefully, no really new issues arise in the solution of the differential equations (6), (7), and (10) for *q*_0_(*k,t*), *q*_1_ (*k,t*), and *S*(*k,t*), respectively. The differential equations are discretized using 4th order BDF with initial time steps using the implicit midpoint rule. The differential equations for *q*_0_(*k,t*), *q*_1_ (*k,t*), and *S*(*k,t*) are solved simultaneously. Therefore, the memory requirements of this algorithm are very low.

The differential equations are solved only for *k* ∈ [*k_min_*(*t*),*k_max_*(*t*)]. During the time step from *t* to *t* + *h*, *k_min_* or *k_max_* (or both) may increase by 1. If *q*_0_ is assumed to be 1 and *q*_1_, *S* are assumed to be zero outside the lens, as stated in the main text, no special handling is necessary when *k_max_* increases. When *k_min_* increases by 1, the values of *q*_0_(*k_min_,t*) as well as *q*_0_(*k_min_*, ·) at previous epochs that contribute to the step to *t* + *h* are taken to be 1. Similarly, the corresponding values of *q*_1_ and *S* are taken to be 0.

### Choice of time step and accuracy

Suppose *dx*/*dt* = –*α*(*t*)*x*. Then

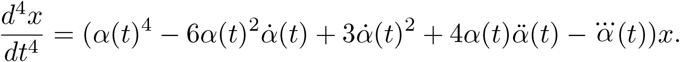

Because the numerical discretizations are formally of order 3, the time step *h* is obtained from the requirement

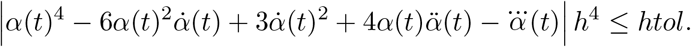

In this requirement, we have taken *x* = 1 because all the differential equations are solving for probabilities. To preserve the numerical stability of the BDF formula, each new time step must be within a factor of 1.2 of the previous time step.

In the computation of the ancestral lens, we take

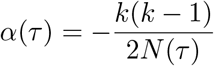

with *k* = *k_max_*(*τ*). In the computation of *q*_0_(*k,t*), *q*_1_(*k,t*), and *S*(*k,t*), we take

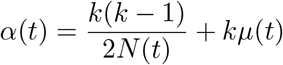

with *k* = *k_max_*(*t*). Derivatives of *α* are computed using differences.

The accuracy of our program has been checked by comparing with implementations of Polanski-Kimmel algorithm [Bhaskar et al., 2014, 2015] and against coalescent simulations [Hudson, 2002]. In addition, we wrote an independent program in Python that solves the differential equations for *k* = 1,…,*n* without limiting itself to an ancestral lens. The accuracy of the C program has been checked against the Python program. The Python program used `odeint()`, which is defined in the `scipy` library. All the reported computations have at least 4 digits of accuracy and often more than 10 digits of accuracy.

### Zero limit with varying mutation rate

Our algorithm to compute the SFS simplifies slightly if we take *μ*(*t*) = *ϵν*(*t*) and then take *ϵ* → 0. The probability *q*_0_(*k,t*) that *k* ancestral samples at time *t* coalesce without being hit by a mutation is then 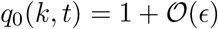. In the limit *ϵ* → 0, we have *q*_0_(*k,t*) ≡ 1.

If we replace *q*_1_(*k,t*) by *ϵq*_1_(*k,t*) in (7) and take *ϵ* → 0, we obtain

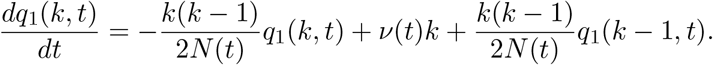

Equations (8), (9), and (10) may still be used to compute the SFS, the only change being the replacement of *μ*(*t*) in (8) by *ν*(*t*).

### The Wright-Fisher SFS with varying mutation rate and population size

Suppose the population size in the Wright-Fisher model is *N_t_* at time *t* and suppose the mutation rate to be *μ_t_* when the *t*-th generation begets the *t* + 1st generation. The probability that *k* samples at time *t* + 1 have *j* parents is given by

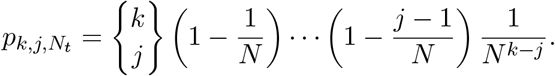

Here 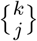 is a Stirling number of the second kind.

Suppose *q*_0_(*k,t*) is the probability that *k* samples in generation *t* coalesce without being hit by a mutation. Then

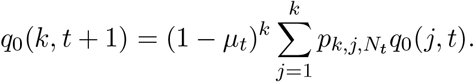

Suppose *q*_1_(*k,t*) is the probability that *k* samples in generation *t* coalesce with exactly one mutation in their Wright-Fisher genealogy. Then

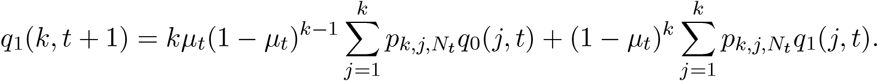

Suppose 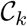 is the condition or event that *k* samples coalesce with exactly one mutation in their Wright-Fisher genealogy. Under the condition 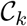, the probability that a sample of size *k* is hit with a mutation during generation *t* + 1 is

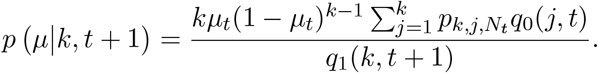

Similarly, the conditional probability that it has *j* parents in generation *t* but does not experience a mutation during generation *t* + 1 is

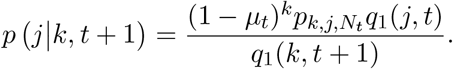

The recurrence for the SFS is then given by

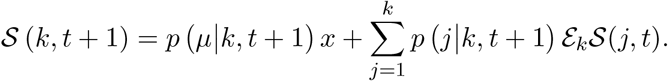

Here 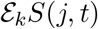 is an extension of the SFS for *j* samples to *k* samples, assuming the *k* samples to be children of the *j* samples. A formula for 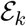 follows from the Wright-Fisher propagators derived in [Melfi and Viswanath, 2018b].

The Stirling numbers are the main source of difficulty in implementing the Wright-Fisher model. However, uniform asymptotic formulas [Temme, 1993] can be used to make the Wright-Fisher implementation much more efficient. The coalescent is not much more than an asymptotic approximation to Stirling numbers of the second kind. If *k* children must be divided between *j* parents, under Wright-Fisher, the children are first partitioned into *j* sets in one of 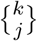 ways and each partition is assigned to a parent. The coalescent produces a partition of *k* children between *j* parents using binary mergers. If *k* – *j* = 1, the two partition distributions are the same. The approximation is not a particularly good one in general, when *k* – *j* ≫ 1, although it works quite well for the SFS [Melfi and Viswanath, 2018b].

